# Fast recurrent processing via ventral prefrontal cortex is needed by the primate ventral stream for robust core visual object recognition

**DOI:** 10.1101/2020.05.10.086959

**Authors:** Kohitij Kar, James J DiCarlo

**Author notes:** correspondence should be addressed to Kohitij Kar.

## Abstract

Distributed neural population spiking patterns in macaque inferior temporal (IT) cortex that support core visual object recognition require additional time to develop for specific (“late-solved”) images suggesting the necessity of recurrent processing in these computations. Which brain circuit motifs are most responsible for computing and transmitting these putative recurrent signals to IT? To test whether the ventral prefrontal cortex (vPFC) is a critical recurrent circuit node in this system, here we pharmacologically inactivated parts of the vPFC and simultaneously measured IT population activity, while monkeys performed object discrimination tasks. Our results show that vPFC inactivation deteriorated the quality of the late-phase (>150 ms from image onset) IT population code, along with commensurate, specific behavioral deficits for “late-solved” images. Finally, silencing vPFC caused the monkeys’ IT activity patterns and behavior to become more like those produced by feedforward artificial neural network models of the ventral stream. Together with prior work, these results argue that fast recurrent processing through the vPFC is critical to the production of behaviorally-sufficient object representations in IT.

## Introduction

A goal of visual neuroscience is to identify and model the brain circuitry that seamlessly solves the challenging computational problem of rapid visual object categorization (DiCarlo and Cox, 2007; Riesenhuber and Poggio, 2000; Yamins and DiCarlo, 2016). Previous studies (Freiwald et al., 2009; Hung et al., 2005; Kar et al., 2019; Logothetis and Sheinberg, 1996; Majaj et al., 2015) show that the pattern of neural activity in the primate inferior temporal (IT) cortex can explicitly represent visual object identities. However, current models of core object recognition fall short of fully explaining both primates’ behavioral image by image difficulty patterns (Geirhos et al., 2017; Rajalingham et al., 2018) and they fall short of fully explaining the distributed population activity patterns of IT neurons (Kar et al., 2019).

These models primarily belong to the family of deep convolutional neural networks (DCNN) with predominantly feedforward architectures. More recent models are beginning to implement recurrent architectures (Kubilius et al., 2019; Nayebi et al., 2018; Spoerer et al., 2017) but experimental data to guide their development is needed. Toward that goal, we have recently demonstrated (Kar et al., 2019) the critical role of putative recurrent signals available at the late-phases of the image evoked IT responses in enabling accurate core object recognition, at least for some images. That study also speculated that the lack of recurrent computations in the feedforward DCNN models might have led to its poor behavioral accuracy and poorer prediction of the late-phase IT responses. But which recurrent circuit motifs in the primate brain are most critical? within ventral stream? Within IT? Top-down from regions downstream of IT (PFC, Amygdala, etc.)? All of the above? Identifying these circuits and inferring their computational functions is critical in developing the next generation of models of the primate visual intelligence and behaviors such as core object recognition.

Kar et al. (2019) determined, for each tested image, the time when response patterns of the IT neuronal population could sufficiently account for the monkey’s object recognition performance on that image, referred to as the object solution time (OST; one OST computed per image). They also identified hundreds of images that critically relied on the early (90-120 ms) and late (150-180 ms) phases of the IT responses post image onset (Figure 1A). These results point to a targeted disruption strategy that we executed here for testing the aforementioned critical recurrent circuits. Specifically, if a particular recurrent circuit motif is critical in core object recognition, its disengagement should: 1) prevent the emergence linearly-decodable object identity information in the late-phase of the IT responses, with little or no effect on the early phase. And, 2) result in a reduction in behavioral performance for the late-solved images, with little or no effect on behavior performance for the early-solved images. In this study, we tested those two predictions for a circuit motif that is recurrently connected to the ventral visual stream — the ventral prefrontal cortex (vPFC).

**Figure 1.**
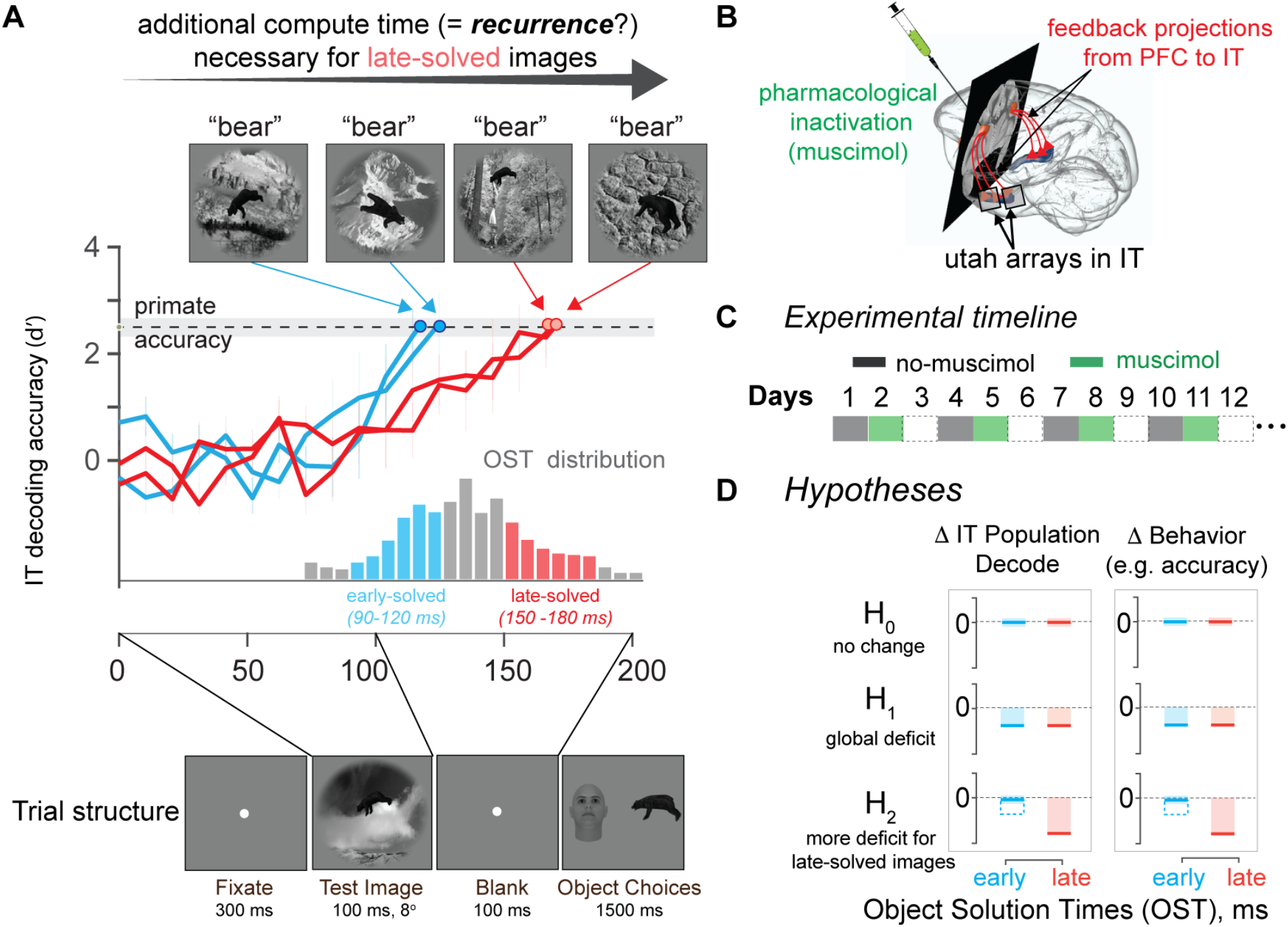
Motivation and Hypotheses. **A**. Temporal evolution of linearly decodable object identity information in IT on an image-by-image basis. For each tested image, we measured the IT population response vector (n=424 neural sites) across time (10 ms resolution). For each time point, we estimated the linear decodable information (cross-validated across images). Each image achieved an IT solution goodness (linear decode accuracy for object identity: d’) which matches the monkey’s behavioral accuracy (avg. of d’=2.5 for the example images, shown as a gray shaded line) after different amounts of processing time (object solution time, OST; gray histogram over; 1320 tested images). Using a range of controls, Kar et al. (2019) concluded that images which exhibit longer OSTs (late-solved; red curves shows two examples) likely require more recurrent processing (relative to images that exhibit shorter OSTs; early solved images; blue curves shows two examples). **B.** Pharmacological inactivation of PFC (ipsilateral to IT recording location) with simultaneous IT population recording. **C.** We divided the experiments into two different sessions, without (gray boxes) and with (green boxes) muscimol injections, conducted on consecutive days. We repeated each session in the same order after a minimum gap of one day (empty boxes). We completed at least 10 sessions of each condition type. **D**. Hypothesized effects of PFC inactivation. One hypothesis (H_0_) is that the robustness of the IT object codes for core object recognition (~200 ms of processing) does not rely at all on PFC, which predicts no change in IT decodes or behavior for both early (blue bar) and late-solved (red bar) images. Another hypothesis (H_1_) is that PFC plays an overall modulatory role in ventral stream computations, which predicts deficits in IT population decode accuracies and behavior that are equal for both groups of images. Finally, a third hypothesis (H_2_) is that PFC as a critical recurrence node in the brain circuitry for core objection, which predicts larger IT population decoding deficits and larger behavioral deficits for late-solved images. A mixture of H_1_ and H_2_ is also possible (see open blue bars; also see Discussion for alternative interpretations).

Among the multiple downstream targets of IT, we chose to first test vPFC because: 1) it is downstream of IT, but has strong recurrent anatomical connections to IT (Borra et al., 2010; Webster et al., 1994; Yeterian et al., 2012), 2) following object category learning, it has been shown to contain object-category selective neurons(Freedman et al., 2001, 2003), 3) previous studies have demonstrated changes in IT resulting from lesion-based (Tomita et al., 1999), pharmacological (Monosov et al., 2011) and thermal perturbation (Fuster et al., 1985) of PFC, and 4) methods to silence PFC are experimentally straightforward because PFC is downstream of IT. Specifically, we here pharmacologically silenced (via muscimol, a GABA^A^ agonist) ~0.4 cm^3^ of ventral PFC in each of two monkeys, and measured changes in IT population activity at the multi-unit level (with chronically implanted Utah arrays ipsilateral to the targeted vPFC, see Figure 1B) and the corresponding changes in core object recognition performance.

Our results show that the inactivation of vPFC reduced the quality of the late-phase IT population activity, as assessed by linear decodability of object identities. We also observed corresponding behavioral deficits during core object recognition tasks — the deficits were significantly higher for late-solved images. Interestingly, the inactivation of vPFC caused the late-phase IT neural activity to become better explained by feedforward DCNN models of the ventral stream. These results argue that fast recurrent processing through the ventral PFC is critical to the production of fully robust object representation in IT and the core object recognition behavior that it supports and that current computational models of the ventral stream lack these computations.

## Results

As outlined above, we reasoned that, if recurrent processing via the ventral PFC to the primate ventral stream is critical for robust core object recognition, then inactivating parts of ventral PFC should produce specific changes in the IT population activity patterns and specific behavioral deficits. In particular, the neural and behavioral deficits should be higher for “late-solved” images — images that we have previously found not to produce a fully formed IT population representation until 150-180 ms post stimulus onset (Kar et al. 2019; see Introduction).

To test the role of vPFC we used pharmacological inactivation of sub-regions of ventral PFC, as previously anatomically landmarked, (Freedman et al., 2003; McKee et al., 2014; Tomita et al., 1999), and identified in this study by structural MRI (see Methods). Based on the expected locations of object category-selective vPFC neurons (Freedman et al., 2001, 2003), we first performed a single electrode measurement survey (Figure S1,C-E) to locate vPFC sub-regions that exhibited strong visual drive and coarse category selectivity (see Methods). We then performed a second structural MRI (now with markers inserted at these locations) to ensure that the localized object-category selective vPFC sites were anatomically consistent with previous reports (Freedman et al., 2001, 2003).

### An assay for recurrent-dependent computation: early-solved vs. late-solved images

Previous studies (Hung et al., 2005; Majaj et al., 2015) have demonstrated that object identity is linearly expressed in the pattern of IT neural activity. Using linear decoders, we have previously estimated the precise time it takes for the macaque IT population to temporally evolve to this linearly explicit pattern for each of 1320 images (Kar et al., 2019; briefly illustrated in Figure 1A). We refer to this time as the object solution time (OST). OST is an estimate (done per image) of the amount of time needed to compute a behaviorally sufficient neural population solution in IT. Longer OSTs, therefore, suggest additional, putatively recurrent computations, beyond what could be achieved by the early, feedforward IT responses. In this study, our analyses primarily focus on comparing the neural and behavioral effects of vPFC inactivation on the images that are solved quickly (“early-solved” images, OST range: 90-120 ms) with the effects on images that are solved slightly later (“late-solved” images, OST range: 150-180 ms).

### vPFC inactivation reduces IT late-phase population activity

We first explored the effect of vPFC inactivation on the quality of the IT neural population patterns evoked by each image. Upon visual inspection (Figure 2B), we observed that vPFC inactivation did not produce a reduction in the (mean) initial (90-120ms) image-driven activity. However, vPFC inactivation appeared to moderately reduce the later portion of the IT responses (i.e., starting around 140 ms after image onset). To look more closely, we compared IT responses at two specific time bins: early phase (90-120 ms; Figure 2C) and late phase (150-180 ms; Figure 2D). We found that, across the entire recorded IT population (n=153 sites), vPFC inactivation produced no significant difference in the mean response (averaged over all images) in the early phase (∆R^early^ = −18 ± 46.4 %, mean ± s.e.m; paired t-test; t(152) = 0.5885, p = 0.5571). However, vPFC inactivation produced a significant reduction in mean late-phase (150-180 ms) IT responses (averaged across images; ∆R^late^ = −31.83 ± 10.4%, mean ± s.e.m; paired t-test; t(152) = 8.5906, p <0.0001). Also, we noted that the time of the emergence of a drop in the mean IT response (black vs. green line in Figure 2B) coincided with the latencies of the vPFC neurons that we recorded at the targeted injection sites (refer Figure S1E) as well as previously measured latencies of neurons in this area (Freedman et al., 2001, 2003). We note that these mean firing rate effects are also consistent with prior causal perturbation studies in other pairs of visually-driven cortical areas (see Discussion).

**Figure 2.**
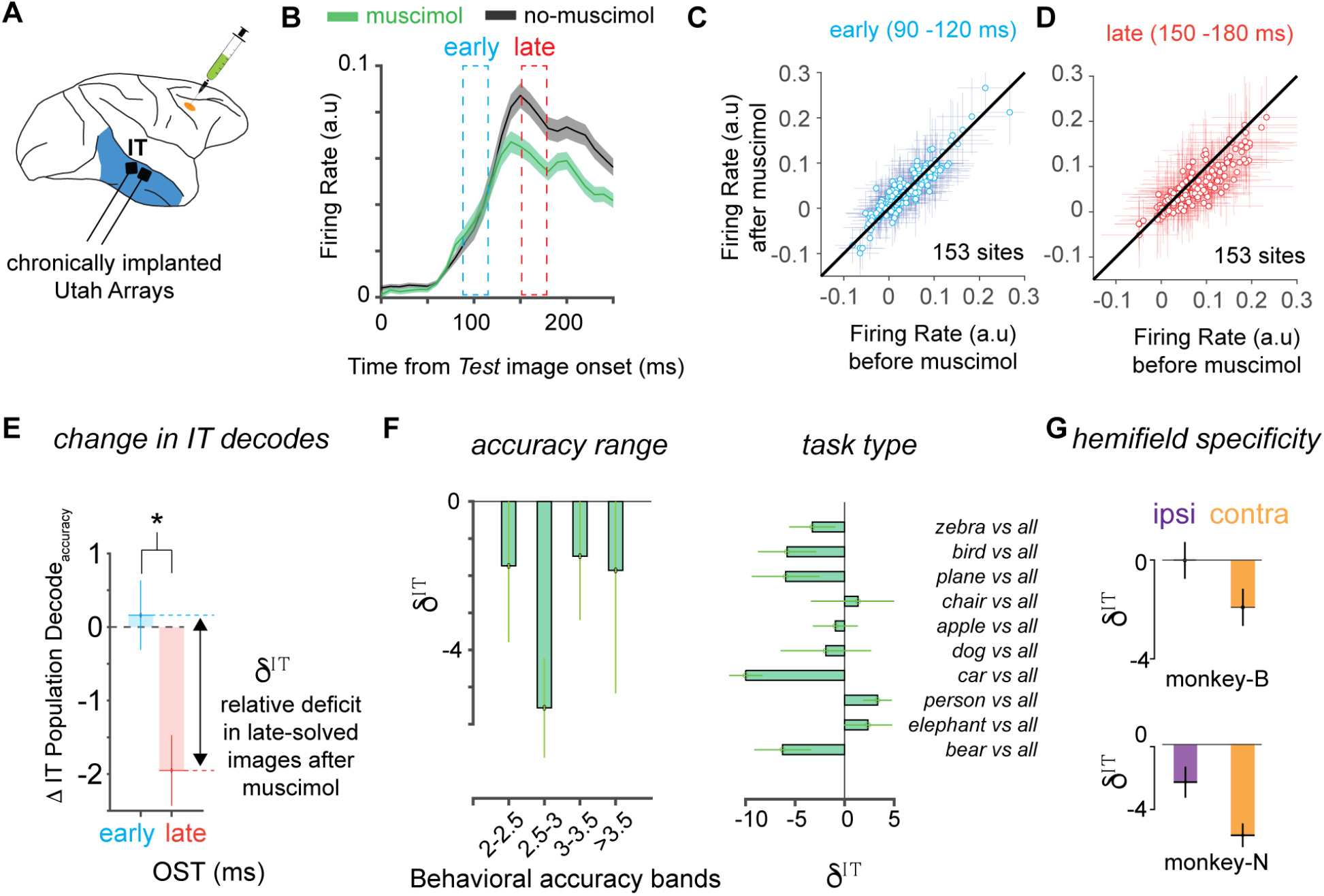
Neural experiments and results. **A.** We measured neural responses from 153 sites in the IT cortex across two monkeys while they performed a battery of core recognition tasks, with and without muscimol injections in the ventral PFC (see Fig. 1C). **B.** Normalized mean IT firing rate in the two conditions (Black = no-muscimol control condition; green = after muscimol injections in PFC). The shades indicate s.e.m across images. **C**. We observed no significant differences across neurons at the early-phase (90-120 ms) of the IT responses (∆R^early^ = −18 ± 46.4 %, mean ± s.e.m; paired t-test; t(152) = 0.5885, p = 0.5571). **D.** We observed a small, but significant reduction in firing rates at the late-phase (150-180 ms) of the IT responses (∆R^late^ = −31.83 ± 10.4%, mean ± s.e.m; paired t-test; t(152) = 8.5906, p <0.0001). Errorbars for C and D denote the standard deviation of responses across images per neuron. **E.** Images (n=234) with late-OST (red bar) showed a significantly higher drop in IT population decode accuracy (see Results) upon vPFC inactivation, compared to the images (n=208) with early-OST (blue bar). This comparison was made with all images that had a measured (behavioral) d’ between 2 and 4, as measured in separate animals (Kar et al. 2019). Error bars denote s.e.m across images. We quantified the strength of this interaction as the difference in the muscimol induced change and we refer to that measure as *δ*^IT^. **F.** The mean *δ*^IT^ was consistently less than 0 for images selected in different ranges of behavioral accuracies. We also observed a negative trend for most, but not all recognition sub-tasks (t-test, t(9) = 1.9718, p=0.0401). Error bars denotes bootstrap CI (95%). **G.** Interaction strength was significantly stronger when we restricted the measurements to images where the object center was in the contralateral visual filed (monkey N, ipsilateral *δ*^IT^:-2%, contralateral *δ*^IT^:-5.8%, permutation test of difference, p<0.001; monkey B, ipsilateral *δ*^IT^: 0.1%, contralateral *δ*^IT^:-2%, permutation test of difference, p<0.001). Error bars denotes bootstrap CI (95%).

### vPFC inactivation selectively disrupts the late-phase IT population code

The neural representations that enable robust object recognition are more subtle than the mean firing rates analyzed above. Indeed, we previously reported that while many images evoke high mean firing rates in the IT cortex, a linearly-readable solution of the foreground object in those images is not present in that activity and emerges only later after subtle changes in the neuron-by-neuron distributed population code (Kar et al., 2019). Thus, we next aimed to examine the temporal evolution of the quality of the IT population code for early-solved versus late-solved images. Here, we assessed “quality” as the ability of the population code to support a linear readout of object identity for held-out test images (i.e., via cross-validation; See Methods). As outlined in the Introduction, we sought to specifically test the hypothesis (H_2_, Figure 1c; right column) that vPFC feedback to the ventral stream, is particularly critical to the development of late-phase IT object solutions. This hypothesis predicts that vPFC inactivation should induce more significant disruptions in the quality of the IT population code for late-solved images compared to the early-solved images at their corresponding object solution times. To control for the behavioral accuracy levels across images, we sub-selected images (out of the total 1320 tested images) for two groups, early-solved (208 images) and late-solved (234 images), that all had a (pre-muscimol) d’ between 2 and 4 (as measured in an earlier study; Kar et al. 2019).

First, we observed that the quality of IT neural population codes (as estimated by linear decode accuracies of object identity) were significantly less accurate at later time points after vPFC inactivation (>150 ms post image onset; median reduction = −2.44%, t-test, t(441) =5.11, p<0.001; Figure S2). Furthermore, to estimate whether the muscimol induced change in IT linear decodability of objects was dependent on the previously estimated OST values (Kar et al., 2019), we compared the IT decode accuracies for the early and late solved images at their corresponding OSTs (Figure 2E). We refer to this difference (early minus late) in the muscimol induced deficits as *δ*^IT^ (as shown in Figure 2E). We observed that vPFC inactivation disrupts the formation of IT solutions for the late solved images more than it disrupts the formation of IT solutions for the early solved images (∆IT Population Decode_accuracy_^early^ = 0.16% ± 0.53 ; ∆IT Population Decode_accuracy_^late^ =−2% ± 0.61, median ± s.e.m ; *t*-test, *t*(441) = 2.4084, *P* = 0.0165; Figure 2E). Moreover, we found that this effect persisted even with different behavioral accuracy level choices (behavioral levels considered in d’: <2, 2-2.5, 2.5-3, >3 ; corresponding *δ*^IT^ were, - 1.63%, −5.63%, −1.57%, −1.9%; Figure 2F). Also, *δ*^IT^ was significantly less than zero considering each of the ten tested objects (10 tasks, t-test, t(9) = 1.9718, p=0.0401; Figure 2F). We observed that the *δ*^IT^ values, when measured separately for each monkey, were significantly more negative for images where the object center was present in the contralateral hemifield (monkey N, ipsilateral *δ*^IT^:-2.01%, contralateral *δ*^IT^:-5.8%, permutation test of difference, p<0.001; monkey B, ipsilateral *δ*^IT^: 0.1%, contralateral *δ*^IT^:-2%, permutation test of difference, p<0.001; yellow bars; Figure 2G), compared to those in the ipsilateral hemifield (purple bars; Figure 2G). Taken together, our results demonstrate that vPFC inactivation disrupts the formation of IT neural population solutions more strongly for images for which those solutions take longer to develop, consistent with the hypothesis that vPFC is part of the critical recurrent circuitry.

### vPFC inactivation produces larger behavioral deficits for late-solved images

As outlined in the Introduction, we hypothesized that if the inactivation of vPFC (Figure 1D) disrupted behaviorally critical recurrent computations (H_2_), then we should expect to see specific changes in IT population codes, and we should also see specific changes in behavior. In particular, we should observe a more significant muscimol induced behavioral performance deficit for images with late OSTs (H_2_; bottom left panel, Figure 1C). The other possibilities are that we observe no change (H_0_; top right panel, Figure 1D) in behavioral performance across images, or an overall shift in the behavioral performance consistently across images with varied OSTs that might indicate a global shift in arousal (H_1_; middle right panel, Figure 1D).

Identical to Kar et al. (2019), in each image, the primary visible object belonged to one of 10 different object categories (Figure 3A). We divided the data collection into two types of sessions — with and without muscimol injections — conducted on consecutive days. These two session types were repeated in an alternative sequence with at least one day of recovery after each muscimol session (Figure 1C; experimental timeline). This design confounds animal satiety and motivation with the effects of muscimol. However, the visual hemifield bias of our reported effects (see below) argue against it. In each session (day), monkeys performed the following tasks sequentially: a passive fixation task, a binary object discrimination task, a second passive fixation task (see Methods). On the second session (day), after the initial passive fixation task, we injected a total of 10 l of muscimol at five depths (2 l each) separated by 0.5 mm in the previously localized ventral PFC area (see Methods for details). We injected in the left hemisphere of monkey B, and right hemisphere of monkey N. Given that the top-down signals from vPFC (both hemispheres) are known to reach both the left and right inferior temporal cortices (Tomita et al., 1999), we have presented our results after the data was pooled across both monkeys. Nevertheless, individual monkey results were consistent with the pooled results (as shown in Figure 3E).

**Figure 3.**
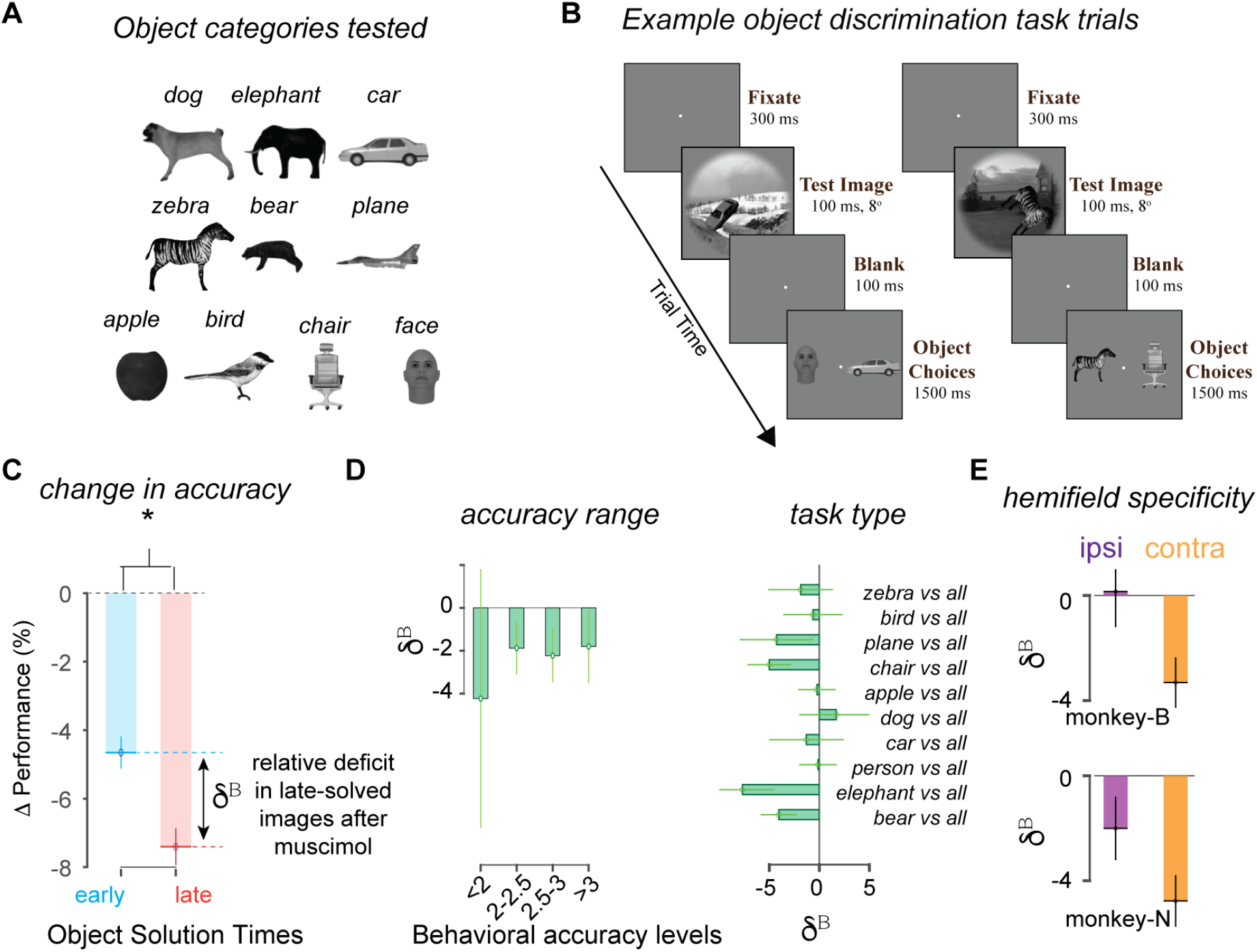
Behavioral experiments and results. **A.** We tested behavioral performance on 10 object categories, where performance was derived from the corresponding 45 binary object discrimination tasks with those 10 categories. **B.** Two example trials of the binary object discrimination task, showing the timeline of events. Monkeys fixate on a central dot, then the test image at 8° containing one of ten possible objects is shown for 100 ms (shown is a car (left trial) and a zebra (right trial)). After a 100-ms delay, a canonical view of the target object and a distractor object (one of the other nine objects) appears (randomly assigned on each trial to the left and right positions), and the monkey indicates which object was present in the test image by making a saccade to one of the two choices. We compared performance on sessions with and without muscimol injections in vPFC **C.** vPFC inactivation resulted in a larger performance drop among images (n=234) with late-OST (red bar), compared to the images (n=208) with early-OST (see Results for statistics). This comparison was made with all images that had a measured d’ between 2 and 4. Error bars denote s.e.m across images. We quantified the strength of this interaction as the difference in the muscimol induced change and we refer to that measure as *δ*^B^. **D.** We observed that the mean *δ*^B^ was consistently less than 0 for images selected in different ranges of behavioral accuracies. We also observed a negative trend for most, but not all recognition sub-tasks (t-test, t(9) = 2.6245, p=0.0276). Error bars denotes bootstrap CI (95%). **E.** We found that the interaction strength was significantly stronger when we restricted the measurements to images where the object center was in the contralateral visual filed (monkey N, ipsilateral *δ*^B^:-2%, contralateral *δ*^B^:-4.8%, permutation test of difference, p<0.001; monkey B, ipsilateral *δ*^B^: 0.1%, contralateral *δ*^B^:-3.3%, permutation test of difference, p<0.001). Error bars denotes bootstrap CI (95%).

First, we observed that there was a significant overall reduction (Performance = 6.03 ± 0.3 %, (mean ± SEM), paired t-test; t(859) = 17.13, p <0.0001; Figure S3A) in performance across all sessions after the muscimol injections. Consistent with hypotheses H_2_ (see Figure 1C), vPFC inactivation caused a significantly higher reduction in performance for late-solved images compared with early solved images (∆Performance^early^ = =−4.76% ± 0.45 ; ∆Performance^late^ =−7.4% ± 0.5, median ± s.e.m ; *t*-test, *t*(441) = 2.3978, *P* = 0.0085). We refer to the difference in the behavioral deficits for the early vs. the late-solved images as *δ*^B^ (as shown in Figure 3C). We observed that *δ*^B^ was consistently negative (i.e., greater behavioral deficits for late-solved images compared to early-solved images) across images grouped according to different behavioral accuracy level choices (behavioral levels considered in d’: <2, 2-2.5, 2.5-3, >3; corresponding *δ*^B^ were, - 4.24%, −1.9%, −2.23%, −1.9%). Also, *δ*^B^ was significantly less than zero considering each of the ten tested objects (10 tasks, t-test, t(9) = 2.6245, p=0.0276). We observed that the *δ*^B^ values when measured separately for each monkey were significantly higher for images in which the object center was in the contralateral hemifield (monkey N, ipsilateral *δ*^B^:-2%, contralateral *δ*^B^:-4.8%, permutation test of difference, p<0.001; monkey B, ipsilateral *δ*^B^: 0.1%, contralateral *δ*^B^:-3.3%, permutation test of difference, p<0.001; yellow bars; Figure 3E), compared to those in the ipsilateral hemifield (purple bars; Figure 3E). We also observed an overall increase in reaction times after muscimol injections (∆RT = 46.3 ± 2.1 ms; t-test; t(858) = 16.3729, p<0.001). Similar to the behavioral accuracy results, we observed that vPFC inactivation increased reaction times, and that this increase was significantly higher for late-solved images than for early-solved images (∆RT^early^ =−34 4.19 ms ; ∆RT^late^ = 55 ± 3.9 ms, median ± s.e.m ; *t*-test, *t*(441) = 2.0488, *P* = 0.04; Figure S3C).

Taken together, these results show that core object discrimination in macaques is disrupted by inactivation of ventral PFC, establishing this area as a critical component of the brain circuitry that is involved in core object recognition.

Furthermore, the performance changes (deficits in accuracy and reaction time) depended on the image being processed — the deficits were more severe for images that more likely depend on recurrent processing (as indexed by each image’s IT object solution time; Kar et al. 2019). These behavioral results are qualitatively consistent with the IT neural results (Figure 2) under the assumption that the behavior is the consequence of mechanisms that are approximated by linear read-out from IT (Majaj et al., 2015).

### Inactivation of vPFC causes the ventral stream to operate more similarly to feedforward computational models

We have previously shown (Kar et al., 2019) that some feedforward deep convolutional neural networks (specific DCNNs) predict the early feedforward (~90-120 ms post image onset) responses of the IT neurons quite well, but are far worse at predicting the late-phase (~150-180 ms) IT responses. These prior results (and other work, see Discussion) suggest that the early-phase IT responses are primarily the product of feedforward computations, but that the late phase IT responses are a more balanced mixture of feedforward and recurrent computation (e.g., through vPFC, as suggested by the results above). Under this hypothesis, the relatively weak ability of these DCNN ventral stream models to explain the late-phase IT responses is due to the lack of the appropriate recurrent computations. If we assume that vPFC inactivation removes those additional recurrent-computations (or blocks the transmission of the results of those computations to IT), vPFC inactivation should make the late-phase IT representations revert to a more feedforward-only mode of operation. vPFC inactivation should, therefore, make the top of the ventral stream operate more like a feedforward only network.

To test this, we used a set of existing feedforward DCNN models (refer Table 1, Methods), and we asked: does vPFC inactivation cause the late-phase IT response to become better explained/predicted by these feedforward models (Figure 4A). We used standard measures of mapping the components of feedforward models onto the responses of individual IT neural sites (Kar et al., 2019; Schrimpf et al., 2018; Yamins et al., 2014; see Methods), and we took the goodness of fit to be the median predictivity across all recorded neural sites. Remarkably, we observed that vPFC inactivation significantly improved the match of the late phase (150-180 ms) IT responses to the feedforward DCNN (AlexNet ‘fc7’) predictions (median late-phase %EV: without muscimol = 21.98%, with muscimol = 28.28%; paired t-test across neurons; t(152)=8.55, p<0.0001; Figure 4B, top panel; also see Figure S4B). Consistent with this, we also found that the vPFC-inactivation caused the late-phase IT responses to be more similar to the early-phase IT responses, as measured by correlation of image response rank order (early vs. late) compared across the muscimol and no-muscimol conditions (paired t-test; t(152)=7.24; p<0.001, see Methods). These results were also consistent across multiple feedforward models (see Figure S4C).

**Table 1.** DCNN model names and corresponding ‘model-IT’ layers used for IT predictivity comparisons (as shown in Figure S4C).

| Model Name | Layer |
| --- | --- |
| AlexNet | fc7 |
| VGG-F | fc7 |
| VGG-S | fc7 |
| SqueezeNet | pool10 |
| VGG16 | fc7 |
| ResNet18 | pool5 |
| VGG19 | fc7 |
| GoogLeNet | pool5-7x7_s1 |
| ResNet50 | avg_pool |
| Inception-v3 | avg_pool |
| ShuffleNet | node_200 |
| Mobilenet_v2 | global_average_pooling2d_1 |
| Xception | avg_pool |
| Resnet101 | pool5 |
| Inception ResNet_v2 | avg_pool |
| DenseNet 201 | avg_pool |

**Figure 4.**
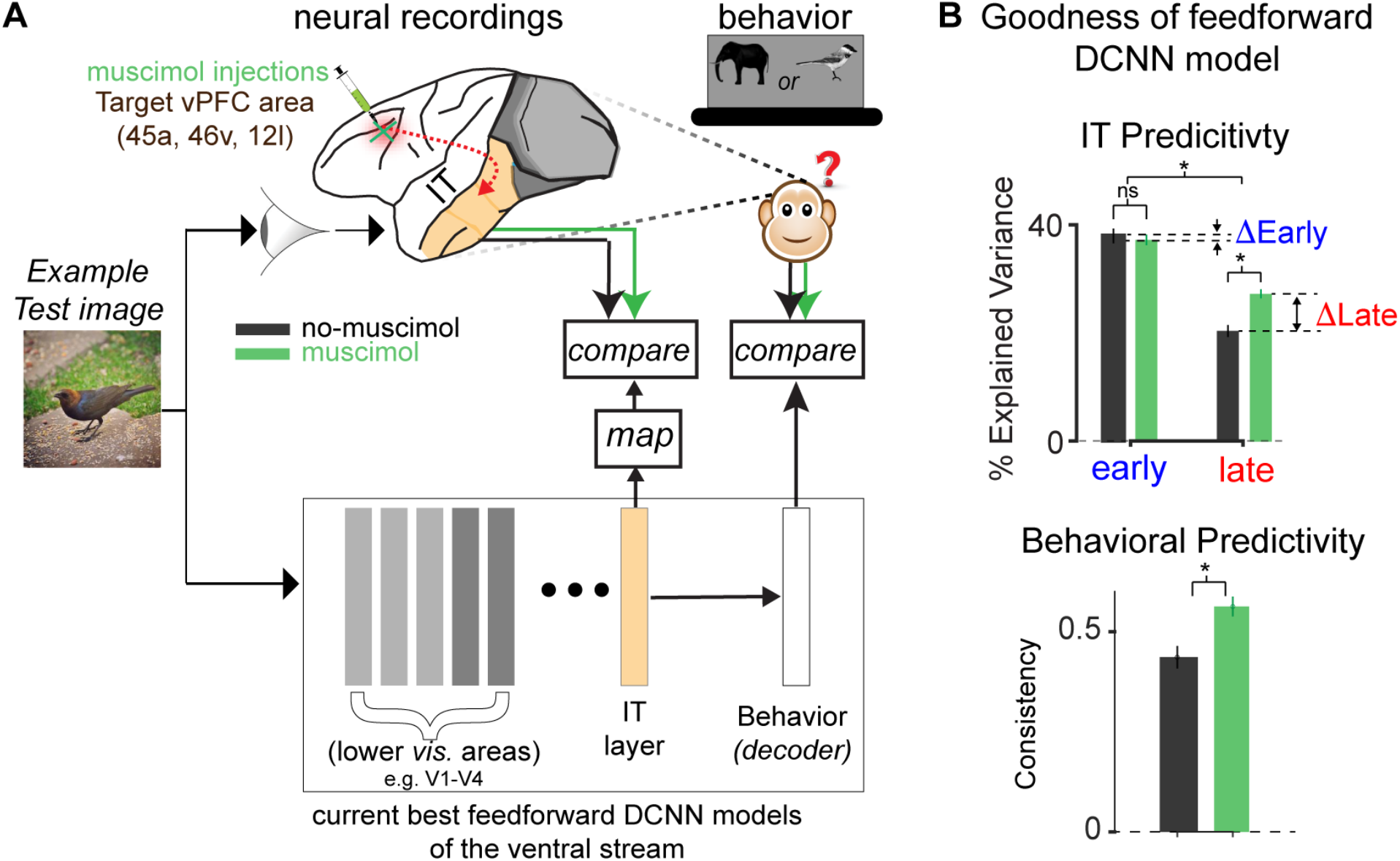
Comparison with computational models: vPFC inactivation causes the ventral stream to behave more like feedforward models. **A.** We showed 683 images to the monkey (fixated passive viewing) while recording simultaneously from their IT cortex, with and without vPFC inactivation (top panel). The dashed red line denotes the recurrently pathway between the ventral PFC and the primate ventral stream. We compared the IT responses with and without vPFC inactivation to those of the penultimate (‘IT’) layers of a feedforward DCNN model of the ventral stream (bottom panel) using previously established methods (see Methods). We also compared the pattern of monkeys’ behavioral responses (pattern of difficulty over images, see Rajalingham et al., 2018) with and without vPFC inactivation to the model’s behavioral pattern. In both types of comparison, the key measure is referred to as predictivity as it assesses the goodness of model predictions on new images. **B.** Top panel: Comparison of IT Predictivity (%EV) of Alexnet (‘fc7’) for early (90-120 ms) and late (150-180 ms) responses, without (black) and with (green) vPFC inactivation. We observed that vPFC inactivation resulted in a significant increase in the match of the late phase of the IT population pattern to the feedforward DCNN “IT” population pattern. No significant changes were observed for early responses. Error bars denote s.e.m across 153 neural sites. Bottom panel: We also observed that vPFC inactivation resulted in a slight, but significant, increase in the match of the monkeys’ object recognition behavior to the object recognition behavior of the feedforward models (match assessed as the correlation (model vs monkey) of the image-by-image pattern of difficulty; see Methods).

We also know from previous work (Rajalingham et al., 2018), that the DCNN models of core object recognition fail to explain primate behavior at an image-by-image level fully— that is, those models do not fully explain and predict which images primates perform well on and which images they perform poorly on. Our results show that vPFC inactivation changed the late phase IT population patterns such that they became better matched to the “IT” layers of feedforward DCNNs. Taken together with prior work that tightly linked primate object recognition behavior to patterns of IT population activity (Majaj et al., 2015), we asked whether the vPFC inactivation also changed the monkey image by image behavior patterns. Indeed, we observed that vPFC inactivation significantly, improved the image by image consistency (normalized correlation, see Methods) between the monkeys’ object recognition behavior and the object recognition behavior of the feedforward models. (Behavioral Predictivity without vPFC inactivation = 0.43, Behavioral Predictivity after vPFC inactivation = 0.56; permutation test of difference; p < 0.0001; Figure 4B right panel).

In sum, these results suggest that vPFC is a critical circuit node that is recurrently modulating the population dynamics of IT. Partially inactivating that node restricts the IT population pattern from correctly evolving away from its initial feedforward response pattern — leaving both the early and the late IT population patterns reasonably well approximated by current feedforward DCNN models of the ventral stream.

## Discussion

In this work, we investigated whether the recurrent circuit connecting the macaque ventral prefrontal cortex (vPFC) to the ventral visual pathway is critical for executing robust core object recognition. We reasoned that if this bidirectional circuitry is indeed critical, then silencing parts of it should produce deficits in the quality of population activity recorded in the IT cortex that is responsible for accurate core recognition behavioral performance. More specifically, based on our prior work (Kar et al. 2019), we hypothesized that we should observe larger deficits for images that take slightly longer to solve and thus their solutions are more likely dependent on recurrent computations (late-solved images; benchmarked earlier in Kar et al., 2019).

Consistent with this hypothesis, we observed that vPFC inactivation produced deteriorations in the quality of the IT population code and deteriorations in behavioral performance that were significantly higher for the late-solved images compared to the early-solved images. Furthermore, we found that vPFC inactivation caused the late phase of the IT population response and the monkey behavior to more closely match the “IT” and behavioral responses of some of the leading feedforward models of the ventral stream. Taken together, these results suggest that vPFC is part of a recurrent circuit that boosts the performance of the ventral stream (relative to shallow feedforward DCNNs) by reshaping the initial (early-phase; putatively feedforward only) neural representations in IT cortex, resulting in corresponding behavioral gains. Consistent with this, removal of vPFC made the ventral stream operate more like a shallow feedforward system. When considered alongside prior work (Kar et al., 2019), this vPFC circuitry is most critical for images that are challenging for shallow feedforward computer vision systems.

### Experimental guidance on developing new scientific hypotheses of ventral stream function

Our current best understanding of neural processing along the ventral stream is carried by specific models in the class of feedforward deep artificial neural networks. These models are the current best scientific hypotheses of the ventral stream because they have the highest overall prediction accuracy (a primary test of a scientific hypothesis: Hempel, 1966; Popper, 1959) for image-evoked responses at all levels of the ventral stream (mean accuracy in V1, V2, V4, IT; Schrimpf et al., 2018). However, because these models do not perfectly predict the image-evoked neural responses of these different areas of the ventral stream (for comparison across different models, see Schrimpf et al., 2018), multiple groups are working to develop even more accurate scientific hypotheses (e.g., Kubilius et al., 2019; Nayebi et al., 2018; Spoerer et al., 2017).

What components do these current models lack? Clearly, the models are missing many things at the single “neuron” level, such as voltage-gated channels to generate spikes, dendritic trees, synaptic components, etc. But we motivated this study by first asking, what critical *network level* components are missing from these models?

Many studies and reviews have suggested the importance of including recurrent circuits to improve such models (Kar et al., 2019; Kietzmann et al., 2019; Lehky and Tanaka, 2016; Tang et al., 2018).This idea is motivated on both anatomical and functional grounds. For example, previous reports (Sugase et al., 1999) have demonstrated that different forms of information can be decoded from early and late responses in IT, suggesting a potential role of intra-areal recurrent inputs to shaping IT population response dynamics. Consistent with the hypothesis that recurrent signals modify late-phase IT population responses, Kar et al. (2019) showed that the ability of feedforward DCNNs to predict the IT population pattern significantly worsened as the IT response pattern evolved. They also showed that this latter portion of the IT population response pattern carries the linearly available object-identity information for many specific images that enable primates to successfully solve them, vastly outperforming shallow feedforward DCNN computer vision models. In sum, the late phase of the IT population response is likely important for robust core recognition behavior, likely depends on recurrent circuits, and it is largely missing from the current best models of the ventral stream. Thus, to produce models of the ventral stream that more closely mimic the mechanisms of the primate brain, a proper form of recurrent network level processing is needed.

But what type of recurrent processing is needed? To begin to answer that question, we here started with an even more basic question: what circuit nodes in the brain are computing and carrying the recurrent signals that we see manifesting as a temporal evolution of the IT late-phase responses? Prior work suggests many potential sources of such signals, including within ventral stream bidirectional pathways, as well as top-down feedback from multiple downstream areas, including vPFC, peri-rhinal cortex, amygdala, and striatum (for review see Kravitz et al., 2013). For reasons outlined in the Introduction, in this study we have specifically focussed on the ventral PFC.

To test the functional importance of a downstream node that is recurrently connected to a target region of interest, many previous studies in the visual system (Bullier et al., 2001; Hupe et al., 1998; Sandell and Schiller, 1982; Wang et al., 2000) have used inactivation methods, similar the one deployed here. In general, those studies report that this downstream manipulation results in a decrease in responses of neurons in the upstream cortical areas, which is analogous to the reduction (~30%) in IT activity level that we have observed here (Figure 2B). For instance, inactivation of area MT (feedback node) via cooling led to a ~20-40% decrease in V1 and V2 responses (refer Figure 1, 2 in Hupe et al., 1998). Focussing specifically on vPFC and IT, prior studies have confirmed that IT responses, similar to other visual areas are modulated by feedback from downstream areas. For example, Fuster et al. (1985) showed that temporary lesions produced by cooling in dorsolateral PFC affected color selectivity in IT neurons. Tomita et al. (1999) performed anterior and posterior commissurectomies, and observed that the responses of IT cortical neurons are modulated by input from the prefrontal cortices, especially for visual information in the contralateral visual field.

Our work is consistent with those studies in that IT responses can be altered by vPFC. However, unlike the work presented here, those earlier studies did not specifically investigate the changes in the distributed IT population code or primate behavior with respect to object recognition, that can guide the development of new models of primate vision. Specifically, that prior work did not engage on questions of the quality of information for recognition behavior at an image by image resolution, or the differential importance of recurrent signals from vPFC as measured in the early vs. late responses of the IT population. Because of this, prior work could not distinguish between an overall modulatory role (H_1_) and a specific set of recurrent computations (similar to H_2_). To our knowledge, the current study is the first to causally test the necessity of the vPFC to ventral stream recurrent circuit at such fast (< 200 ms), but natural time scales, with simultaneous large-scale neural and behavioral measurements. Here, we have leveraged our previous findings (as resported in Kar et al., 2019) to employ a targeted disruption strategy for identifying critical recurrent circuits using pre-defined challenge images (that take additional solution times in IT). Therefore, our results provide evidence that feedback from vPFC does not simply modulate IT (e.g., gain) — it specifically improves the format of the distributed IT population code, and those improvements are specific to the late phase of this code.

However, the results reported here do not identify the exact circuitry involved in the re-entry of information from the vPFC into the ventral stream. Previous anatomical studies have shown that the feedforward projections that connect the ventral stream to the prefrontal cortices originates in the anterior portions of the lower ventral bank and fundus of the STS (for a review, see Kravitz et al., 2013) and mainly target areas 45A/B, 46v, and 12r/l in the ventrolateral prefrontal cortex (VLPFC). On the other hand, feedback projections from these same areas in the PFC are distributed across the inferior temporal cortical areas TEO and TE (Gerbella et al., 2010). There is not much evidence of direct connections between these areas in the PFC and earlier visual areas (V1, V2, and V4). But, we cannot rule out the possibility of indirect connections to the lower visual areas via the frontal eye fields and other regions.

Each of these possible circuit motifs is a hypothesis that must next be implemented as a set of neural network models for future experimental testing. Our neural measurements (with and without vPFC inactivation, as reported here) can be used to select among such models. For instance, we can estimate the weights of the feedback connections between vPFC and the ventral stream nodes such that the model approximates the neural firing rates at its IT layer (as measured here) upon random (~0.4 cm^3^) lesions of the ventral PFC module.

Many studies (Ganis et al., 2007; Harth et al., 1987; Tang et al., 2018) propose a cognitive role of the prefrontal feedback: the idea that these recurrent connections carry an expectation signal that augments the representation of object identity in the IT cortex. Our results are consistent with this and other similar conceptual theories. However, those ideas are not specific enough to be tested for individual images. That is, they do not specific how to build an accurate image-computable neural network model of the IT-to-PFC-to-IT circuit. While the results presented in this study do not provide a precise blueprint for such a model, the temporal and image level specificity that they build on is already useful for guiding the development of new recurrent, image-computable models (Kubilius et al., 2019) and the current results can further guide the placement and simulation testing of a PFC node in such models (see more below).

### Role of vPFC in core object recognition behavior

Previous work (Freiwald et al., 2009; Hung et al., 2005; Kar et al., 2019; Logothetis and Sheinberg, 1996; Majaj et al., 2015) has linked neuronal responses in the inferior temporal cortex to primate core object recognition behavior. For instance, Majaj et al. (2015) experimentally rejected a large number of alternative models that link ventral stream population activity to core object recognition behavior (aka “decoding models” or “linking models”) in support of a simple linear weighted sum of IT response model. These models posit that the mechanisms of core object recognition beyond IT are: approximately linear sums of the activity levels of individual IT neurons computed by neurons in PFC, peri-rhinal cortex, or in the caudate. Using various combinations of model parameters (e.g., numbers of neurons, amount of experience with each object category, brain location of the downstream linear summing), multiple linking hypotheses can be constructed. Our results do not narrow the space of hypotheses to a single linking model. However, these experiments have provide two architectural constraints for new models, and our data can be used to falsify or support each such model. First, vPFC is required to support core object recognition behavior and therefore needs to be integrated into any future model of such behavior. Second, feedback signals conveyed via the recurrent connections between vPFC and the ventral stream (most likely the IT cortex) are also necessary to support this behavior. Below we speculate and discuss candidate linking models that can be further developed and tested using our results.

One possibility is that the downstream summing nodes (as posited by previous studies) are vPFC neurons and those vPFC neurons drive the monkey’s behavior. According to this hypothesis, the vPFC is an additional, bidirectionally connected node of processing that intervenes between IT and behavior. This idea is conceptually simple, and it is motivated by previous data from vPFC, including results showing that category training in monkeys causes PFC neuronal responses to become categorical-like (Freedman et al., 2001), which is what would be expected if vPFC was the location of those learned sums of IT neuronal responses described above. This hypothesis predicts that vPFC inactivation should lead to an *equal* decrease in behavioral performance for every image. An alternate possibility, however, is that vPFC neurons do not drive behavior directly, but they instead transmit the product of their computations to support upstream brain regions, such as the IT cortex, which then drive behavior via other brain nodes such as caudate. This second possibility is also consistent with the prior work that demonstrated category selectivity in vPFC neurons (Freedman et al., 2001). Our data do not unequivocally resolve among these two possibilities. Our results that vPFC inactivation leads to larger deficits for late-solved images (Figure 3C) is consistent with the second possibility. However, the fact that vPFC inactivation also led to significant deficits for early-solved images argues for some element of the first possibility. Indeed, our results overall seem to suggest that both ideas may be partially correct.

Experimentally, we speculate that large-scale neural measurements in brain-regions like vPFC, collected simultaneously in behaving monkeys (solving a wide variety of recognition tasks), will be required to gain further insights. Furthermore, feedback projection-specific causal perturbation experiments (similar to Oguchi et al., 2015) will be necessary to identify and functionally characterize some of these circuit motifs. However, to drive further progress, we now need to incorporate the circuit motifs and build specific artificial neural network models motivated by these experimental results, test their image by image predictions, eliminate models that do not match the experimental data, and build new models. That iterative cycle will ultimately lead to a complete, neurally mechanistic understanding of visual object recognition, from images to behavior.

## Methods

### Subjects

The nonhuman subjects in our experiments were two adult male rhesus monkeys (***Macaca mulatta***).

### Visual stimuli: generation

All stimuli used in this study were previously used in the Kar et al. 2019 study. For a brief description of the stimuli (see Figure S1B for example images), please refer below.

#### Generation of synthetic (“naturalistic”) images

High-quality images of single objects were generated using free ray-tracing software (http://www.povray.org), similar to Majaj et al. (2015) Each image consisted of a 2D projection of a 3D model (purchased from Dosch Design and TurboSquid) added to a random background. The ten objects chosen were bear, elephant, face, apple, car, dog, chair, plane, bird and zebra (Figure 3A). By varying six viewing parameters, we explored three types of identity while preserving object variation, position (*x* and *y*), rotation (*x*, *y*, and *z*), and size. All images were achromatic with a native resolution of 256 × 256 pixels.

#### Generation of natural images(photographs)

Images pertaining to the 10 nouns, were download from http://cocodataset.org. Each image was resized to 256 × 256 × 3 pixel size and presented within the central 8. We used the same images while testing the feedforward DCNNs.

### Primate behavioral testing

#### Active binary object discrimination task

We measured monkey behavior from two male rhesus macaques. Images were presented on a 24-inch LCD monitor (1920 × 1080 at 60 Hz) positioned 42.5 cm in front of the animal. Monkeys were head fixed. Monkeys fixated a white dot (0.2°) for 300 ms to initiate a trial. The trial started with the presentation of a sample image (from a set of 1360 images) for 100 ms. This was followed by a blank gray screen for 100 ms, after which the choice screen was shown containing a standard image of the target object (the correct choice) and a standard image of the distractor object. The monkey was allowed to view freely the choice objects for up to 1500 ms and indicated its final choice by holding fixation over the selected object for 400 ms. Trials were aborted if gaze was not held within ±2° of the central fixation dot during any point until the choice screen was shown.

#### Passive Fixation Task

During the passive viewing task, monkeys fixated a white dot (0.2°) for 300 ms to initiate a trial. We then presented a sequence of 5 to 10 images, each ON for 100 ms followed by a 100 ms gray (background) blank screen. This was followed by fluid reward and an inter trial interval of 500 ms, followed by the next sequence. Trials were aborted if gaze was not held within ±2° of the central fixation dot during any point.

#### Eye Tracking

We monitored eye movements using video eye tracking (SR Research EyeLink 1000). Using operant conditioning and water reward, our 2 subjects were trained to fixate a central white square (0.2°) within a square fixation window that ranged from ±2°. At the start of each behavioral session, monkeys performed an eye-tracking calibration task by making a saccade to a range of spatial targets and maintaining fixation for 500 ms. Calibration was repeated if drift was noticed over the course of the session.

#### Data collection

We divided the data collection into two different sessions (with and without muscimol injections; Figure 1B) conducted on consecutive days. These two sessions were repeated in the same order with a minimum gap of one day post the muscimol session (Figure 1C; experimental timeline). On each session (day), monkeys performed the following tasks sequentially: a passive fixation task, a binary object discrimination task, a second passive fixation task. On the second session (day), after the initial passive fixation task, we injected a total of 10 l of muscimol at 5 depths (2 l each) separated by 0.5 mm in the previously localized ventral PFC area (for details see below).

### Behavioral Metrics

We have used a one-vs-all image level behavioral performance metric (similar to the one used in Kar et al., 2019) to quantify the behavioral performance of the monkeys as well as DCNNs (described below). This metric estimates the overall discriminability of each image containing a specific target object from all other objects (pooling across all 9 possible distractor choices).

Given an image of object ‘*i*’, and all nine distractor objects (*j* ≠ *i*) we computed the average performance per image as,

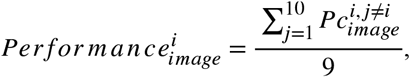

where *P*_*c*_ refers to the fraction of correct responses for the binary task between objects ‘*i*’ and ‘*j*’.

To compute the reliability of this vector, we split the trials per image into two equal halves by resampling without substitution. The median of the Spearman-Brown corrected correlation of the two corresponding vectors (one from each split half), across 1000 repetitions of the resampling was then used as the reliability score (i.e. internal consistency).

### Large scale multi-electrode recordings and simultaneous pharmacological inactivation

#### Surgical implant of chronic micro-electrode arrays

We surgically implanted each monkey with a head post under aseptic conditions. After behavioral training, we recorded neural activity using 10 × 10 micro-electrode arrays (Utah arrays; Blackrock Microsystems). A total of 96 electrodes were connected per array. Each electrode was 1.5 mm long and the distance between adjacent electrodes was 400 μm. Before recording, we implanted each monkey multiple Utah arrays in the IT cortex (monkey B: 2 arrays in left hemisphere); monkey N: 2 arrays in the right hemisphere). Array placements were guided by the sulcus pattern, which was visible during surgery. The electrodes were accessed through a percutaneous connector that allowed simultaneous recording from all 96 electrodes from each array. Behavioral testing was performed using standard operant conditioning (fluid reward), head stabilization, and real-time video eye tracking. All surgical and animal procedures were performed in accordance with National Institutes of Health guidelines and the Massachusetts Institute of Technology Committee on Animal Care.

#### Surgical implant of PFC injection chamber

During the same surgery, as the chronic array implant, we also placed a semi-cylindrical chamber (Crist Instruments) over a craniotomy targeting the prefrontal cortex, around the principal sulcus. We placed the chambers in the left and right hemispheres of monkey B and monkey N respectively. The chambers were held in place by dental acrylic (methyl methacrylate) applied around the chamber. We used previously reported anatomical landmarks (Freedman et al., 2003; McKee et al., 2014; Tomita et al., 1999), identified by an initial MRI, to guide the PFC chamber placements.

#### PFC injection protocol

During the sessions with muscimol injections, we first carefully scraped the dura for maximal visibility and minimum resistance in the path of injection. Then, we used an in-house set up to lower the injection needles (30-32 gauge, small Hub RN Needle; Hamilton Company) using a micro-syringe pump and controller (Micro4™ World Precision Instruments). We started approximately 4 mm below the estimated surface of the dura. We injected 0.5 uL of muscimol (5mg/mL, Sigma Aldrich) at that depth at a speed of 1000 nL/min and, waited for 3 mins and pulled the needle up by ~0.5 mm. This was repeated for 5 depths in total. After the end of the final injection, we waited for 30 mins before the start data collection.

### Electrophysiological Recording

During each recording session, band-pass filtered (0.1 Hz to 10 kHz) neural activity was recorded continuously at a sampling rate of 20 kHz using Intan Recording Controller (Intan Technologies, LLC). The majority of the data presented here were based on multiunit activity. We detected the multiunit spikes after the raw data was collected. A multiunit spike event was defined as the threshold crossing when voltage (falling edge) deviated by more than three times the standard deviation of the raw voltage values. Of 384 implanted electrodes, 2 arrays (left and right hemispheres for monkey B and N respectively) × 96 electrodes × two monkeys, we focused on the 153 most visually driven, and reliable neural sites. Our array placements allowed us to sample neural sites from different parts of IT, along the posterior to anterior axis. However, for all the analyses, we did not consider the specific spatial location of the site, and treated each site as a random sample from a heterogeneous pool of IT neurons.

### Neural recording quality metrics per site

#### Visual drive per neuron 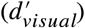

We estimated the overall visual drive for each electrode. This metric was estimated by comparing the image responses of each site to a blank (gray screen) response.

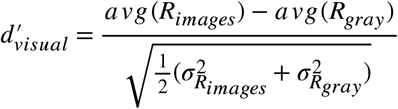

#### Image rank-order response reliability per neural site 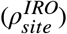

To estimate the reliability of the responses per site, we computed a Spearman-Brown corrected, split half (trial-based) correlation between the rank order of the image responses (all images).

#### Inclusion criterion for neural sites

For our analyses, we only included the neural recording sites that had an overall significant visual drive 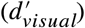, and an image rank order response reliability 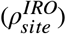 that was greater than 0.6. Given that most of our neural metrics are corrected by the estimated noise at each neural site, the criterion for selection of neural sites is not that critical. It was mostly done to reduce computation time and eliminate noisy recordings.

### Estimation of IT population decode accuracies at OST

To estimate what information downstream neurons could easily “read” from a given IT neural population, we used a simple, biologically plausible linear decoder (i.e., linear classifiers), that has been previously shown to link IT population activity and primate behavior (Majaj et al., 2015). Such decoders are simple in that they can perform binary classifications by computing weighted sums (each weight is analogous to the strength of synapse) of input features and separate the outputs based on a decision boundary (analogous to a neuron’s spiking threshold). Here we have used a support vector machine (SVM) algorithm with linear kernels. The SVM learning model generates a decoder with a decision boundary that is optimized to best separate images of the target object from images of the distractor objects. The optimization is done under a regularization constraint that limits the complexity of the boundary. We used L2 (ridge) regularization, where the objective function for the minimization comprises of an additional term (to reduce model complexity),

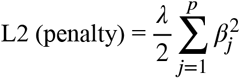

where *β* and p are the classifier weights associated with ‘p’ predictors (neurons). A stochastic gradient descent solver was used to estimate 10 (one for each object) one-vs-all classifiers. After training each of these classifiers with a set of 100 training images per object, we generated a class score (*sc*) per classifier for all held out test images given by,

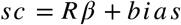

where R is the population response vector and the bias is estimated by the SVM solver. The train and test sets were pseudo-randomly chosen multiple times until we every image of our image set was part of the held-out test set. Only the responses from the no-muscimol conditions were treated as training signal. All predictions were made either on held-out responses from no-muscimol or muscimol conditions. We then converted the class scores into probabilities by passing them through a *softmax* (normalized exponential) function.

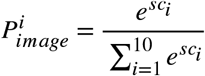

In our previous study (Kar et al., 2019), object solution time per image, *OST*_*image*_was defined as the time it takes for linear IT population decodes to reach within the error margins of the pooled monkey behavioral accuracy for that image. Given that we have used the exact same images in this study, we have used our previously estimated OST per image as the time point of comparison of IT decode accuracy, 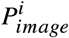 (with and without muscimol) per image. All reported values of IT population decode accuracies are estimates of how well the population decode accuracy was at the specific OST estimated for the specific image.

### Estimating change in image-driven response rank order (early vs late)

For each neuron we estimated the image response vector 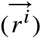 at two specific time bins (early: 90-120 ms, and late: 150-180 ms; post image onset). To estimate the change in this vector across time, we computed the noise corrected correlation between the 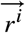 vectors estimated at the early and late time bins respectively, as follows,

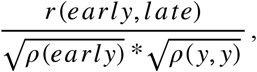

where r(early,late) is the correlation between the 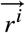 vectors estimated at the early and late time bins, and *ρ*(early) and *ρ*(late) are the split-half (across trial) reliability of these vectors estimated independently at the corresponding time bins. We computed these noise-corrected correlation values per neuron for both the no-muscimol and muscimol conditions.

### Binary object discrimination tasks with DCNNs

We have used the same linear decoding scheme mentioned above (for the IT neurons) to estimate the object solution strengths per image for the DCNNs. Briefly, we first obtained an imagenet pre-trained DCNN (AlexNet). We then replaced the last three layers (i.e., anything beyond ‘fc7’) of this network with a fully connected layer containing 10 nodes (each representing one of the 10 objects we have used in this study). We then trained this last layer with a back-end classifier (L2 regularized linear classifier; similar to the one mentioned for IT) on a subset of images from our image-set. These images were selected randomly from our imageset and used as the train-set. The remaining images were then used for the testing (such that there is no overlap between the train and test images). Repeating this procedure multiple times allowed us to use all images as test images providing us with the performance of the model for each image.

### Prediction of neural responses from Deep Convolutional Neural Networks (DCNN) features

We modeled each IT neural site as a linear combination of the DCNN model features. We first extracted the features per image, from the DCNNs’ penultimate layers. Using a 10-fold train/test split of the images, we then estimated the regression weights (i.e., how we can linearly combine the model features to predict the neural site’s responses) using a partial least squares (MATLAB command: *plsregress*) regression procedure, using 20 retained components. For each set of regression weights estimated on a train imageset, we generated the output of that ‘synthetic neuron’ for the held out test set. The percentage of explained variance, *IT predictivity* (for more details refer Yamins et al., 2014) for that neural site, was then computed by normalizing the r^2^ prediction value for that site by the self-consistency of the image responses for that site and the self-consistency of the regression weights (similar to Kar et al. 2019) for that site (estimated by a Spearman Brown corrected trial-split correlation score). Table 1 lists all the models we have tested and the corresponding layers treated as “model-IT”.

## Data and code availability

At the time of publishing the data associated with all the figures and the code used to generate the figures will be available upon reasonable request.

## Acknowledgements

This research was supported by the Office of Naval Research MURI-114407 (J.J.D), and by the Center for Brains, Minds and Machines (CBMM), funded by NSF STC award CCF-1231216. We thank K.M. Schmidt, A.R. Murthy and S. Sanghavi for technical assistance.

## Author contributions

K.K. and J.J.D designed the experiments and data analyses pipelines. K.K. carried out the experiments. K.K. performed the data analyses. K.K. and J.J.D. wrote the manuscript.

---

**Figure S1.**
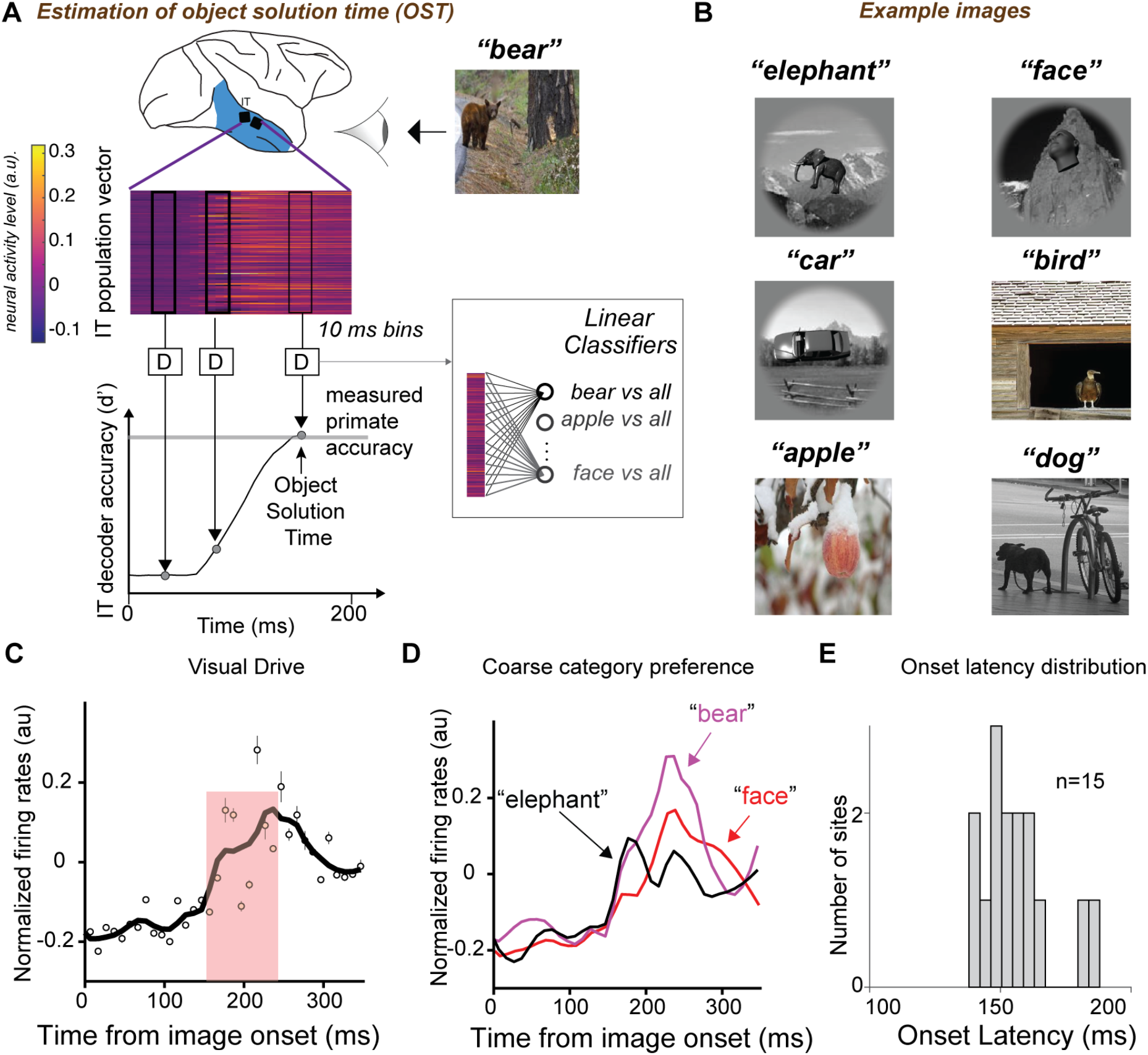
A. Estimation of object solution time. For each image (example image of a bear shown) presentation (100 ms), we counted multiunit spike events (see Methods for details), per site, in non overlapping 10 ms windows, post stimulus onset to construct a single population activity vector per time bin. These population vectors (image evoked neural features) were then used to train and test cross-validated linear support vector machine decoders (D) separately per time bin. The decoder outputs per image (over time) were then used to perform a binary match to sample task, and obtain neural decode accuracies at each time bin. The time at which the neural decodes equal the primate (monkey) performance, is then recorded as the object solution time (OST) for that specific image. B. Six example images from six different object categories. C. Sample neural response from a vPFC site (averaged across 10 repetitions and 80 images). D. Coarse category selectivity of an example vPFC neuron. Each curve is the average response per object category (8 images per category, 10 repetitions per image). E. Distribution of onset latencies of 15 neural sites in vPFC (7 in monkey B, 8 in monkey N).

**Figure S2.**
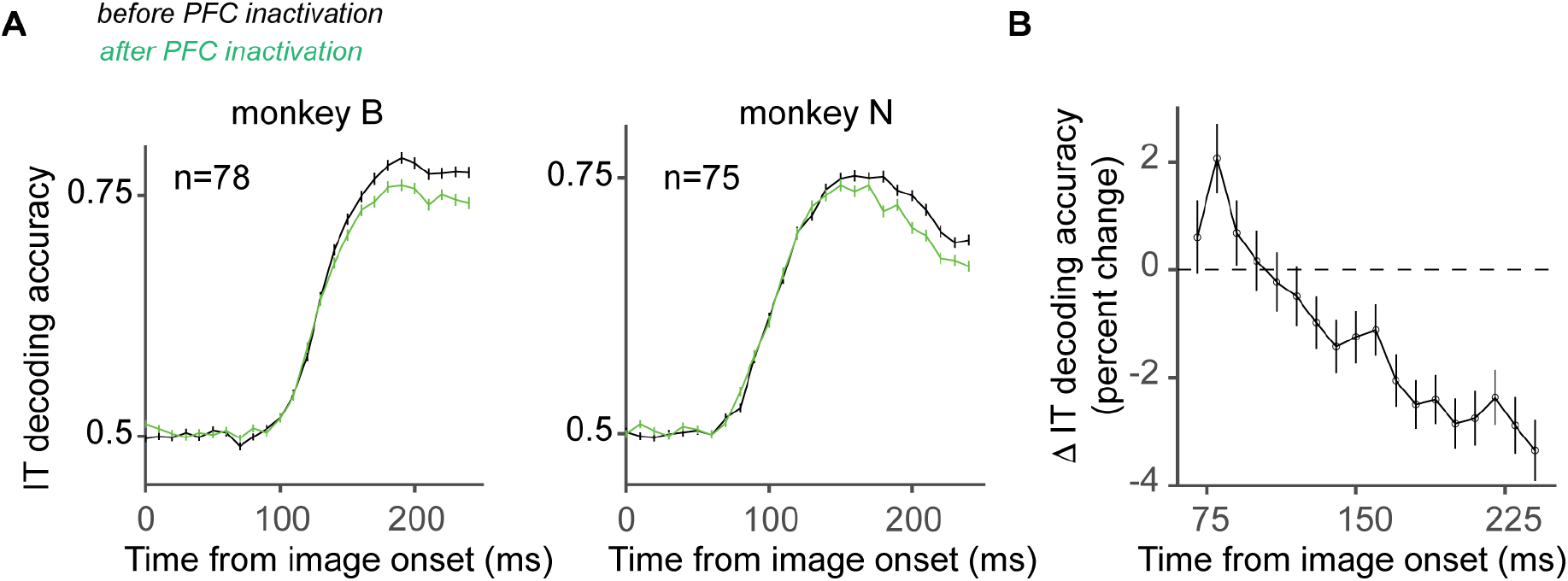
A. IT population decoding accuracies (percent correct) estimated independently for each monkey; left panel: monkey B (78 sites); right panel: monkey N (75 sites) with (green) and without (black) vPFC inactivation. We observe that in each monkey, the quality of the IT population code drops after vPFC inactivation ~150 ms post image onset. B. Dependence of the drop in quality of IT population code, i.e., difference between the green and the black curves shown on the left; estimated on the pooled neural population (153 sites). Error-bars denote s.e.m across images.

**Figure S3.**
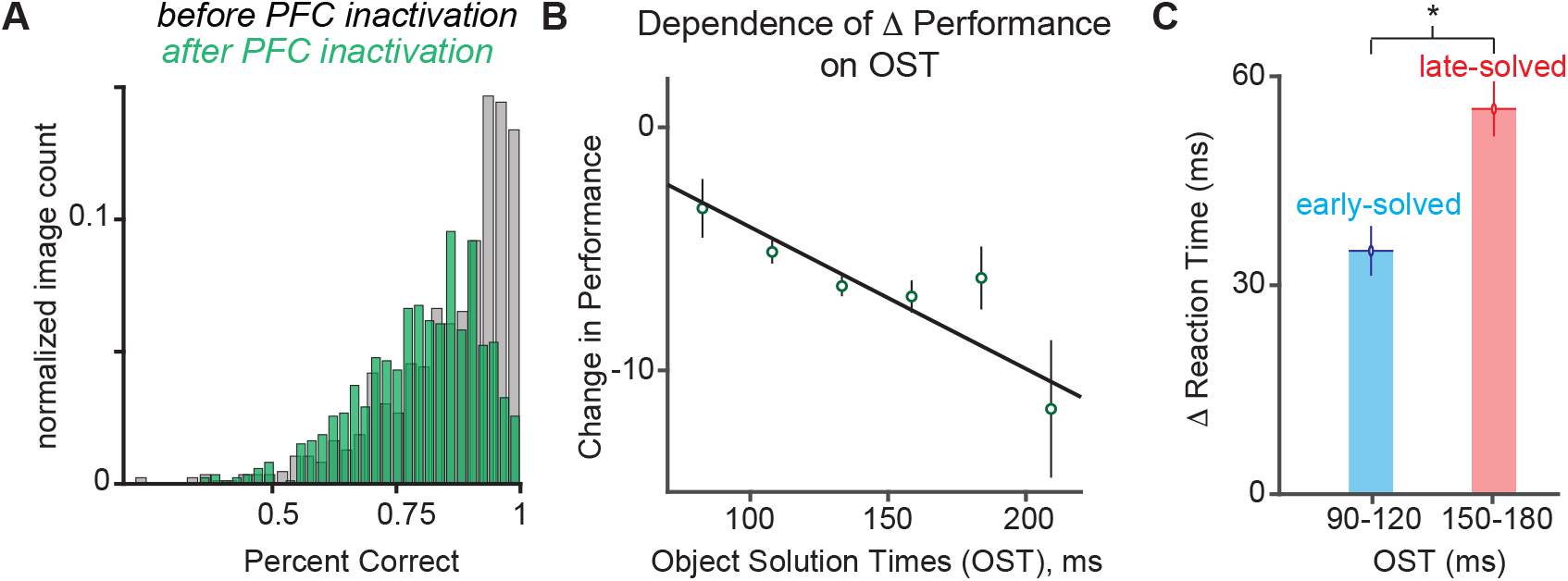
A. Distribution of behavioral performances (percent correct) with (green) and without (black) vPFC inactivation. There was a significant overall reduction (∆Performance = 6.03 ± 0.3 %, (mean ± SEM), paired t-test; t(859) = 17.13, p <0.0001) in performance across all sessions after the muscimol injections. B. ∆ Performance across different bins of object solution times. ∆Performance was negatively correlated with object solution times (Spearman R = −0.11; p = 0.0015; computed image by image). TheError-bars denote s.e.m across images. C. vPFC inactivation increased reaction times, and that this increase was significantly higher for late-solved (red) images than for early-solved (blue) images (∆RT^early^ =−34 ± 4.19 ms ; ∆RT^late^ = 55 ± 3.9 ms, median ± s.e.m ; *t*-test, *t*(441) = 2.0488, *P* = 0.04). Error-bars denote s.e.m across images.

**Figure S4.**
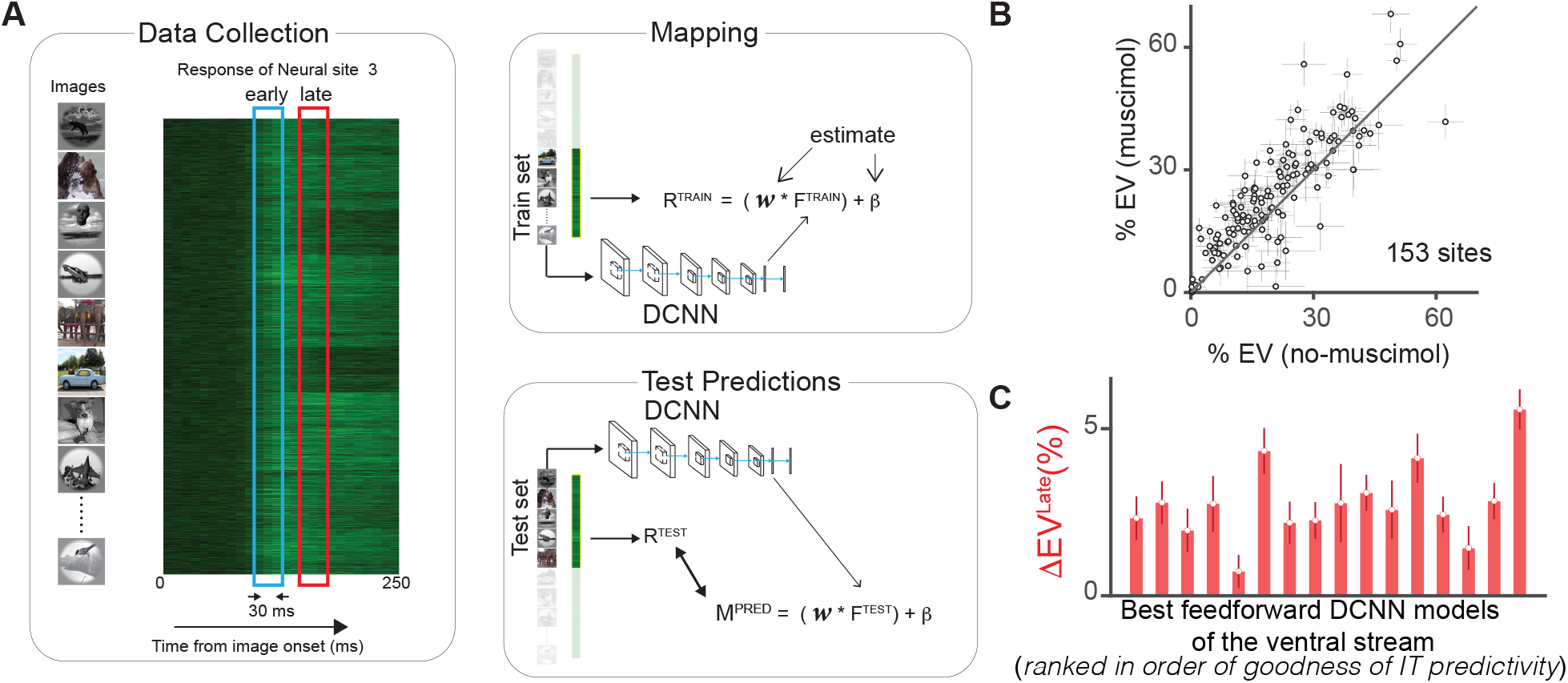
A. Predicting IT neural responses with DCNN features. Schematic of the DCNN neural fitting and prediction testing procedure. This includes three main steps. Data collection: neural responses are collected for each image e.g. example neural site is shown, across 10 ms timebins Late and early time window is demarcated. Mapping: We divide the images and the corresponding neural features (R^TRAIN^) into a 50-50 train-test split (shown for demonstration). For the train images, we compute the image evoked activations (F^TRAIN^) of the DCNN model from a specific layer. We then use partial least square regression to estimate the set of weights (*w*) and biases (*β*) that allows us to best predict R^TRAIN^ from F^TRAIN^. Test Predictions: Once we have the best set of weights (*w*) and biases (*β*) that linearly map the model features onto the neural responses, we generate the predictions (M^PRED^) from this synthetic neuron for the test image evoked activations of the model F^TEST^. We then compare these predictions with the held-out test image evoked neural features (R^TEST^) to compute the IT predictivity of the model. B. Comparison of how well (in units of percentage of explained variance) Alexnet (‘fc7’) features predict the neural responses (153 sites across 2 monkeys) measured with and without PFC inactivation. % EV estimated post muscimol injections were significantly higher than ones estimated without vPFC inactivation. Error-bars denote standard deviation across cross-validation split repetitions. C. We observed that the difference in %EV (with and without vPFC inactivation) for the late-phase (150 - 180 ms) IT responses was significantly greater (muscimol > no-muscimol) than 0, for all 15 tested feedforward networks (plotted on the x-axis; ranked in order of goodness of IT predictivity). Error bars denote s.e.m across 153 neural sites.

